# Time resolved multi-omics reveals diverse metabolic strategies of *Salmonella* during diet-induced inflammation

**DOI:** 10.1101/2024.02.03.578763

**Authors:** Katherine Kokkinias, Anice Sabag-Daigle, Yongseok Kim, Ikaia Leleiwi, Michael Shaffer, Richard Kevorkian, Rebecca A. Daly, Vicki H. Wysocki, Mikayla A. Borton, Brian M. M. Ahmer, Kelly C. Wrighton

**Affiliations:** Department of Microbiology, Immunology, and Pathology, Colorado State University, Fort Collins, Colorado, USA; Department of Microbial Infection and Immunity, The Ohio State University, Columbus, Ohio, USA; Department of Chemistry and Biochemistry, The Ohio State University, Columbus, Ohio, USA; Department of Cell and Molecular Biology, Colorado State University, Fort Collins, Colorado, USA; Department of Soil and Crop Sciences, Colorado State University, Fort Collins, Colorado, USA

**Keywords:** RNA-seq, time series, respiration, microbial metabolism, pathogenesis, CBA/J mice

## Abstract

With a rise in antibiotic resistance and chronic infection, the metabolic response of *Salmonella enterica* serovar Typhimurium to various dietary conditions over time remains an understudied avenue for novel, targeted therapeutics. Elucidating how enteric pathogens respond to dietary variation not only helps us decipher the metabolic strategies leveraged for expansion but also assists in proposing targets for therapeutic interventions. Here, we use a multi-omics approach to identify the metabolic response of *Salmonella enterica* serovar Typhimurium in mice on both a fibrous diet and high-fat diet over time. When comparing *Salmonella* gene expression between diets, we found a preferential use of respiratory electron acceptors consistent with increased inflammation of the high-fat diet mice. Looking at the high-fat diet over the course of infection, we noticed heterogeneity of samples based on *Salmonella* ribosomal activity, which separated into three infection phases: early, peak, and late. We identified key respiratory, carbon, and pathogenesis gene expression descriptive of each phase. Surprisingly, we identified genes associated with host-cell entry expressed throughout infection, suggesting sub-populations of *Salmonella* or stress-induced dysregulation. Collectively, these results highlight not only the sensitivity of *Salmonella* to its environment but also identify phase-specific genes that may be used as therapeutic targets to reduce infection.

**Importance:** Identifying novel therapeutic strategies for *Salmonella* infection that occur in relevant diets and over time is needed with the rise of antibiotic resistance and global shifts towards Western diets that are high in fat and low in fiber. Mice on a high-fat diet are more inflamed compared to those on a fibrous diet, creating an environment that results in more favorable energy generation for *Salmonella*. Over time on a high-fat diet, we observed differential gene expression across infection phases. Together, these findings reveal the metabolic tuning of *Salmonella* to dietary and temporal perturbations. Research like this, exploring the dimensions of pathogen metabolic plasticity, can pave the way for rationally designed strategies to control disease.

## Introduction

*Salmonella enterica* serovar Typhimurium (*Salmonella*) is a leading cause of gastrointestinal disease worldwide, posing serious public health risk due to increasing antibiotic resistance (1, 2). One challenge of controlling this pathogen is its broad metabolic capacity and adaptability to its environment. Recent studies have demonstrated that *Salmonella* infection can be modified with a robust microbiome and through diet manipulation(3–7). However, mechanisms explaining these diet-based phenomena remain understudied. Here we address this knowledge gap by leveraging deeply sequenced, time-series transcriptomics, to reveal the metabolism of *Salmonella* throughout infection.

As a facultative anaerobe, *Salmonella* outcompetes the native microbiota through initiation of the host’s inflammatory response and numerous virulence factors(8–11). In addition to oxygen diffusion into the gut lumen, the subsequent inflammatory response results in the generation of reactive oxygen and nitrogen species which produce respiratory electron acceptors such as nitrate, nitrite, dimethyl sulfoxide (DMSO), trimethylamine N-oxide (TMAO), fumarate, tetrathionate and thiosulfate(12–18).

Furthermore, various carbon sources become available to *Salmonella* with inflammation. *Salmonella* can utilize host and microbial metabolic end products, such as lactate and ethanolamine, as well as microbial derived succinate, and more energetically favorable carbon sources(18–24). Most studies evaluating *Salmonella* substrate and electron acceptor use focus on a single compound. Moreover, these studies often do not track metabolism under different dietary conditions or over time, primarily focusing on late-stage infection processes.

Diet is a critical driver of gut microbiomes, influencing the gut metabolic landscape and microbial membership, which can alter colonization resistance against *Salmonella*(25–28). For example, high-fat diets (HFDs) result in increased inflammation and host susceptibility to infection(29–31). Furthermore, prior research demonstrated that pre-treatment with a high-fat, low-fiber, Western diet was sufficient to break pathogen colonization resistance, resulting in increased susceptibility to *Salmonella*(3). Given the expansion of the Western diet globally(32), studying *Salmonella* pathogenesis and physiology in more realistic diet backgrounds is needed.

Changes in the microbial membership, chemical landscape of the gut, and host response are dynamic factors exploited by pathogens like *Salmonella*(9, 11, 13). Yet, time-series studies are limited in this field. Here we investigate *Salmonella* metabolic processes by analyzing gene expression from CBA mice fed fibrous or high-fat diets and over time. Pairing 16S rRNA sequencing, metatranscriptomic sequencing, lipocalin-2 analysis, and both targeted and untargeted metabolomics, we revealed known and previously unrecognized metabolic strategies that distinguish early, peak, and late infection phases. These data emphasize the importance of environmental context to *Salmonella* metabolism and demonstrate preferential expression of metabolic and pathogenic pathways by diet and infection phase. These key pathways could be targeted to abate enteric infection.

## Results and Discussion

### High-fat diet increases inflammation and *Salmonella* respiratory electron acceptor utilization

Using fecal samples, we assessed the role of diet on *Salmonella* infection by comparing the effects of a fibrous chow diet (Chow) or a high-fat diet (HFD) on *Salmonella* relative abundance, *Salmonella* gene expression, and mouse inflammation (**Fig. 1A**). First, we used 16S rRNA amplicon sequencing (16S) to screen the relative abundance of *Salmonella* per mouse and compared microbial community metrics between the two diets. We assessed nine *Salmonella* infected mice on days 8 (HFD) and 11 (Chow), along with paired pre-infection samples from the same mice (n=18). Days 8 (HFD) and 11 (Chow) were chosen as late infection samples based on disease severity(33), *Salmonella* relative abundance, and sample availability (see methods).

**Figure 1:**
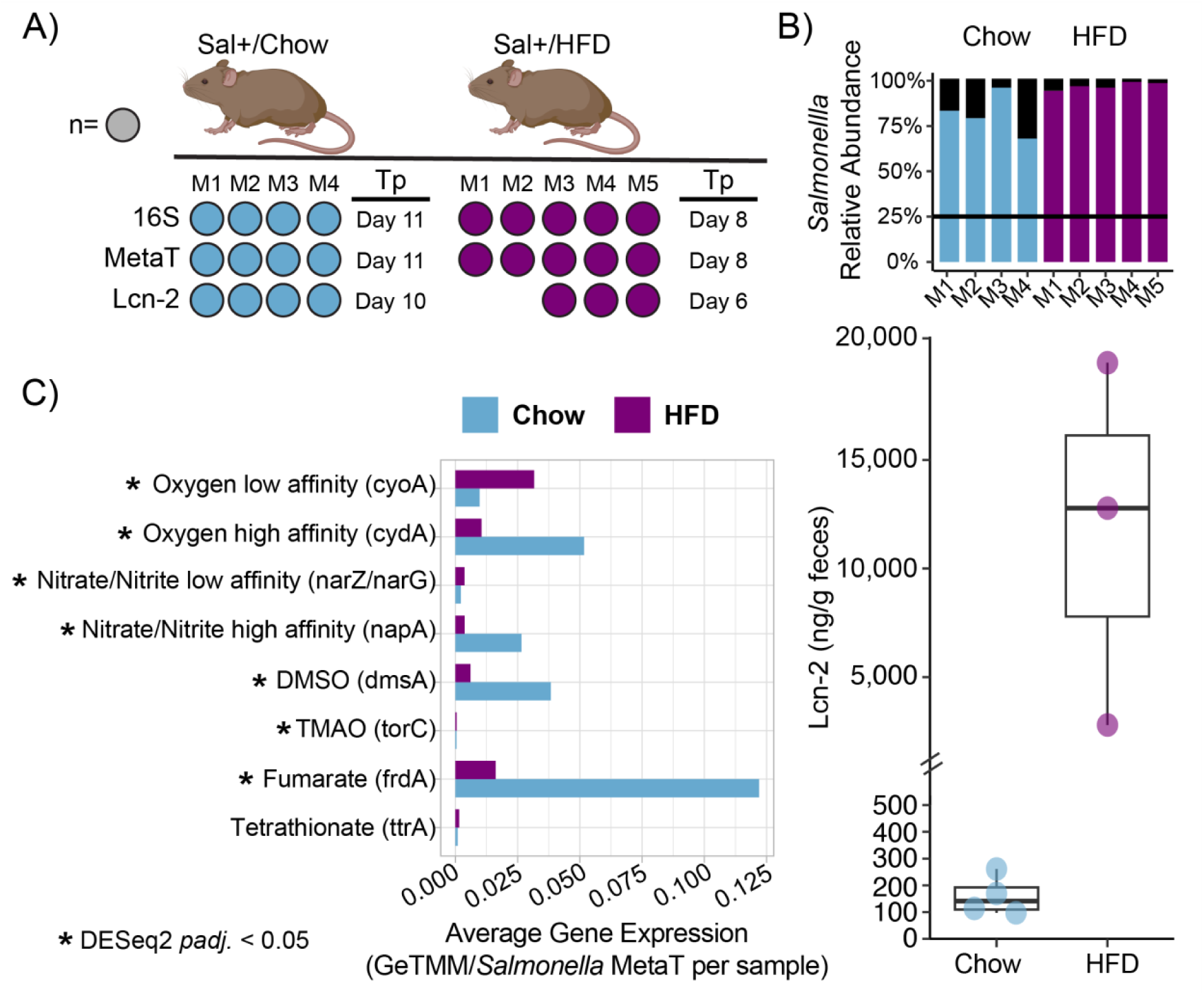
Increased inflammation and use of respiratory electron acceptors when comparing HFD and Chow mice. A) Experimental design figure describes the number of mice, the dietary treatment, and types of analysis that were conducted. 16S rRNA amplicon sequencing (16S), metatranscriptomics (MetaT), and lipocalin-2 ELISA assays (Lcn-2) were conducted on fecal samples from mice fed either chow diet (Chow, blue) or high-fat diet (HFD, purple) with timepoint (Tp) indicated. B) Box plot shows the median and Q1/Q3 -/+ 1.5 interquartile range of concentrations of Lipocalin-2 (ng/g), as a measure of inflammation, from fecal samples for Chow and HFD mice. Stacked bar chart shows *Salmonella* abundance (colored blue or purple based on diet) for each mouse relative to the rest of the microbial community (denoted by black bars) as determined from 16S rRNA amplicon sequencing. Black line denotes samples that are high responders (25% *Salmonella* relative abundance). C) Bar chart shows average *Salmonella* gene expression as gene length corrected trimmed mean of M-values (GeTMM) for key respiratory electron acceptors between Chow and HFD mice. Asterisk (*) indicates that the gene was significantly expressed between dietary regimes by DESeq2 (*adjusted p* value <0.05).

All selected mice, regardless of diet, were classified as high responders with *Salmonella* relative abundance >25% (**Fig. 1B, top**)(9, 33). The HFD (97%) had a slightly higher average *Salmonella* relative abundance compared to the Chow (82%). Despite these slight differences, there was no significant difference in microbial diversity between the two treatments at late infection (ANOVA, *p*=0.386 and *p*=0.133) (**Fig. S1**). Regardless of dietary treatment, the paired pre-infection samples exhibited decreased microbial richness and Shannon’s diversity compared to their respective post-infection samples. Consistent with reports from others(3, 34), Chow pre-infection samples had significantly higher microbial diversity than HFD pre-infection samples (ANOVA, *p*<0.001 and *p*=0.008, respectively). This finding suggests that diet disrupts the microbiome, potentially impacting *Salmonella* physiology.

Along with *Salmonella* relative abundance, we measured lipocalin (Lcn-2) concentrations, which is a host-derived protein indicating inflammatory status(35). Lcn-2 concentrations (ng/g of feces) illustrated a significant increase in inflammation in HFD mice compared to Chow mice (**Fig. 1B**, bottom) (ANOVA, *p*<0.001). Together, our 16S and lipocalin analyses illustrate that while diet alone can reduce microbial diversity, the presence of *Salmonella* results in more pronounced inflammation in HFD mice during late infection.

To ensure higher fidelity of our experimental results and address potential strain heterogeneity that may have developed during laboratory maintenance of this strain(36–39), we constructed a draft genome for this *Salmonella* isolate. This pangenome was derived from a combination of short and long read sequencing (see methods). This strain-resolved genome shared 4597 called genes with 99.99% average nucleotide identity to the previously published *Salmonella* ATCC genome (SAMN08777876). Our sampling averaged 27.19 Gbp of sequencing per sample, generating 1,363,165,050 reads. The internally derived genome was used to map metatranscriptomic sequences from nine fecal samples (Chow=4, HFD=5) and resulted in consistent read mapping regardless of diet.

Prior reports have suggested that inflammation increases electron acceptor availability(8, 11, 12, 20), which favors *Salmonella* growth during infection. As such, we hypothesized that we would see increased respiratory electron acceptor expression concurrent with increased inflammation in the HFD. *Salmonella* gene expression revealed that oxygen (*cyoA* and *cydA*), nitrate and nitrite (*narZ*/*narG*, and *napA*), DMSO (*dmsA*), TMAO (*torC*), and fumarate (*frdA*) utilization genes were differentially expressed in the HFD compared to the Chow diet **(Fig. 1C, Data set S2)**. Tetrathionate reduction (*ttrA*), while detected, did not show significant expression differences across diet treatments.

Additionally, when *Salmonella* encodes multiple genes for utilizing an electron acceptor, like oxygen and nitrogen, we observed increased expression of genes that function optimally at higher substrate concentrations and are more energetically favorable. Specifically, in the HFD, *Salmonella* preferentially expressed the low-affinity oxygen (*cyoA*) and nitrate (*narZ/narG*) utilization genes, compared to the less inflamed Chow where *Salmonella* activated the high-affinity oxygen (*cydA*) and nitrate (*napA*) utilization genes(40–44). Collectively, our results indicate that *Salmonella* exploits HFD-induced inflammation and suggests that *Salmonella* can finely tune its energetic strategy to local chemical conditions.

### *Salmonella* respiration is structured by infection phase

Given the elevated inflammation and metabolic response in HFD mice, we were interested in observing the progression of *Salmonella* metabolism throughout infection. We collected fecal samples from five HFD mice 1, 2, 3 and 6 days before *Salmonella* inoculation, and continued daily sampling after inoculation until sacrifice (day 8). Fecal samples were processed for 16S rRNA amplicon sequencing (16S), metatranscriptomics (metaT), lipocalin-2 ELISA assays (Lcn-2), targeted short-chain fatty acids (SCFAs) metabolomics, and untargeted metabolomics (LC-MS) (**Fig. 2A**).

**Figure 2:**
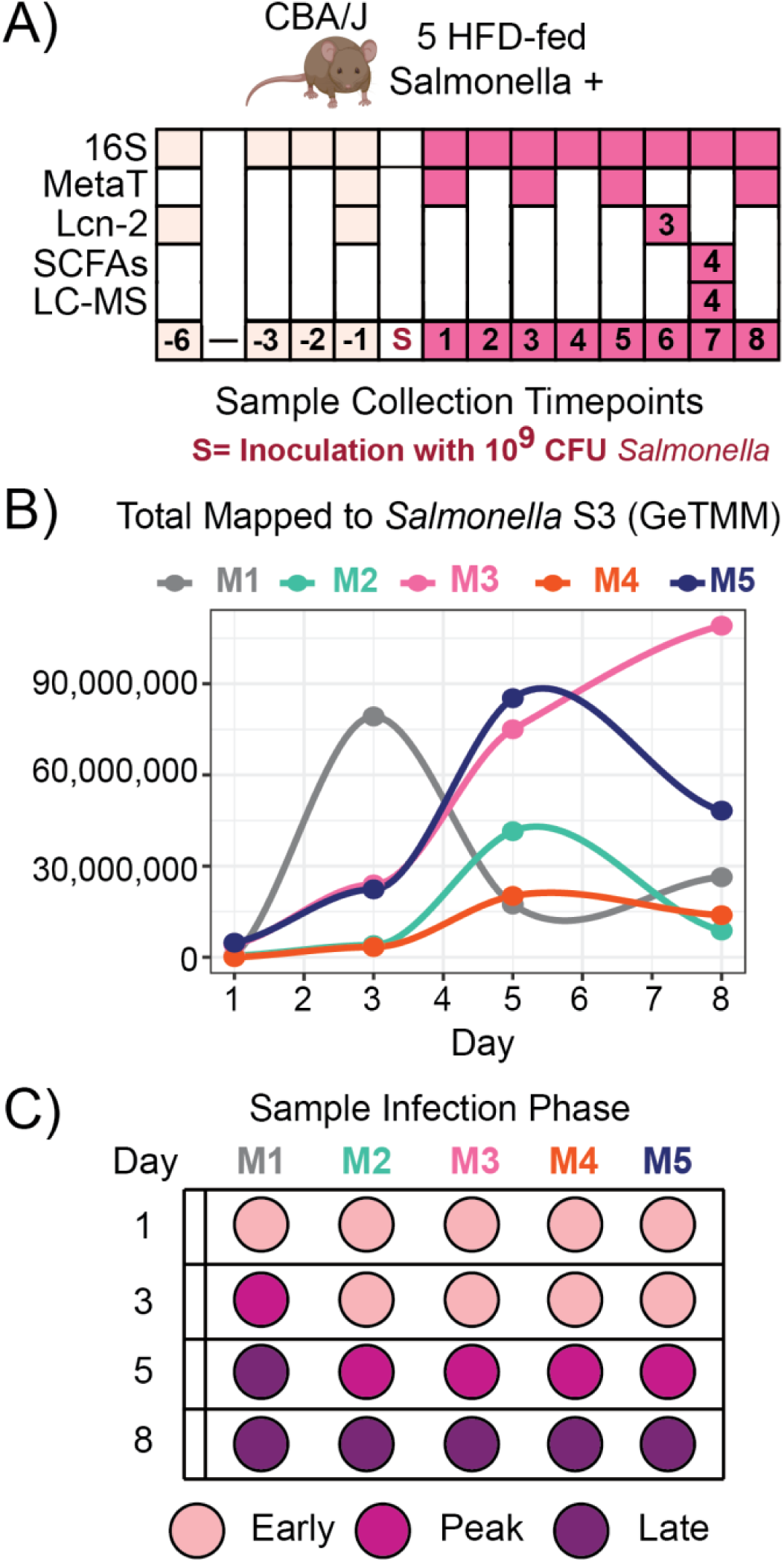
*Salmonella* infection heterogeneity over time. A) High-fat diet (HFD) experimental design of 5 mice describes the types of analysis that were conducted on fecal samples over time. 16S rRNA amplicon sequencing (16S), metatranscriptomics (MetaT), lipocalin-2 ELISA assay (Lcn-2), targeted short-chain fatty acids (SCFAs) metabolomics, and untargeted metabolomics (LC-MS) were conducted prior to infection (light pink) and after inoculation with 10^9^ CFU *Salmonella* (dark pink). Analyses were performed from fecal samples from all 5 mice, unless noted otherwise by numbers within colored boxes. B) The line plot shows normalized single-copy marker gene expression (GeTMM) of the *Salmonella* S3 ribosomal protein per mouse over time. C) Given the heterogeneity of *Salmonella* gene expression over time, samples were grouped into infection phase (early: light pink, peak: dark pink, and late:purple).

Amplicon sequencing was used to profile *Salmonella* relative abundance across all 60 samples, guiding metatranscriptomic sample selection (**Data set S1**). We collected metatranscriptomes during infection days 1, 3, 5, and 8, as well as day −1, analyzing 5,339,114,584 reads from 25 samples, with an average depth of 32.25 Gbp per sample (**Data set S2**).

Timeseries amplicon data showed that all HFD mice became high responders by day 5, but we note that there was heterogeneity among mice in the timing of peak *Salmonella* relative abundance (**Fig. S2**). Consequently, we used expression of the single-copy S3 ribosomal protein (*rpsC*) from *Salmonella* to group samples by infection phase relative to each mouse over time (**Fig. 2B**). As shown in Figure 2B, peak expression of *Salmonella* varied over time and between mice. Using the relative increase of S3 gene expression per mouse, we clustered the samples into 3 infection phases: early (9 samples), peak, (5 samples), and late (6 samples) (**Fig. 2C**) (see methods).

Using our metatranscriptomics data and the sample grouping described above, we compared expression of oxygen (*cyoABCDE* and *cydAB*), nitrate (*narGVZ*), TMAO (*torA*), tetrathionate (*ttrS*), thiosulfate (*phsABC*) and hydrogen (*hybC*) utilization genes (**Fig. 3**). These genes were differentially expressed between the infection phases according to DESeq2 or clustering by GeTMM normalization (**Data set S2**). Our findings revealed selective utilization of respiratory electron acceptors during infection phases (**Fig. 3**). Early and late infection phases exhibited increased expression of anaerobic respiration genes (*narGVZ*, *phsABC*), while peak phase showed increased expression of aerobic respiration genes (*cyoABCDE* and *cydAB)*. Of these respiratory complexes, only the catalytic subunits for tetrathionate showed no differential temporal signal; however, the sensor for tetrathionate (*ttrS*) was distinctive in the early samples. Consistent with prior research, this respiratory capacity provides *Salmonella* a competitive advantage against obligate fermentative microorganisms prevalent in the pre-infection gut(8, 33, 45–47).

**Figure 3:**
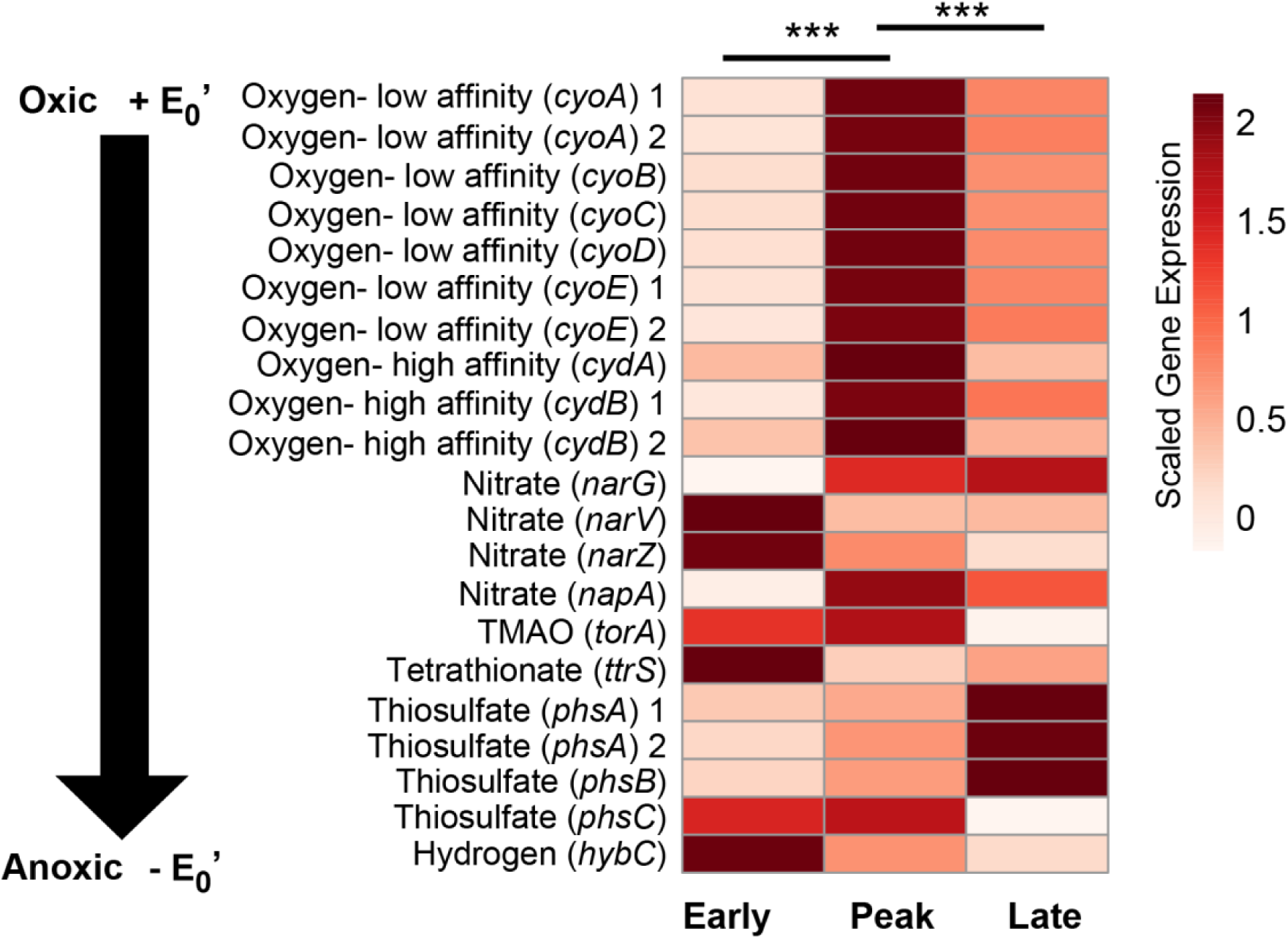
Respiratory electron acceptors utilization by infection phase. Heatmap of the mean, normalized gene expression from respiratory electron acceptor utilization genes mapped to the *Salmonella* pangenome shows patterns between infection phases. Genes are listed from oxic to anoxic (black arrow), and 15 samples are grouped by infection phase (early, peak, and late). Asterisks indicate statistical significance between phases where *** is a *p* value of <0.001.

Our findings indicate differential electron acceptor use along the infection gradient. It was not surprising to see oxygen use, the most energetically favorable electron acceptor, at peak infection when *Salmonella* ribosomal protein expression was also highest, as the lumen becomes more oxygenated in response to *Salmonella*(13). Interestingly, genes encoding anaerobic respiration were more highly expressed in the early and late infection phases. These data also demonstrate that multiple electron acceptor genes are activated simultaneously in the same infection phase, possibly reflecting sub-population responses across the gut habitats(38, 48, 49), or co-metabolic regulatory control under common redox transcriptional regulators(50, 51), findings warranting further investigation. This dynamic gene expression highlights how an energetically versatile bacterium like *Salmonella* rapidly optimizes its energetic strategies to changing local chemical conditions during infection and as a consequence of host-pathogen-commensal microbiota interactions(23, 52–54).

### Targeted and untargeted substrate profiles revealed during infection

Prior studies have reported the multitude of electron donors *Salmonella* can competitively utilize during respiration. In some cases, it is thought that the pathogen utilizes lower-energy carbon substrates not viable for commensal obligate fermenters. Some of these include ethanolamine and 1,2-propanediol, which have been suggested to be important for *Salmonella* expansion over commensal microbes(18, 20, 22, 55). Additionally, higher energy carbon sources, such as mannitol, arabinose, and galactitol, have been studied in relation to intracellular survival, *Salmonella* expansion, or competition(4, 6, 23, 56).

While tracking the expression of genes that utilize these carbon sources throughout different infection phases, we noticed significant expression changes for galactitol (*gatD*) during early infection. Notably, mannitol (*mtlA*, *mtlD*) and arabinose (*araA*, *araB*, *araD*) were expressed across all infection phases and did not uniformly show enrichment over any infection phase. Additionally, ethanolamine (*eutC*) and 1,2-propanediol (*pduC*) were not discriminate of a particular infection phase but instead predominately expressed during both the peak and late phases. In summary, our study design allowed us to track gene expression of substrate use over time, adding new insights into *Salmonella* occupancy throughout infection.

Furthermore, our untargeted approach provided the potential to discover new putative substrates that may support *Salmonella* expansion, especially in this less-explored high-fat diet model. To do this, we examined the global clustering of substrate related gene expression, comparing it with our metabolite data from HFD infected, HFD uninfected, and Chow uninfected mice (**Data set S3, Data Set S4**). The genes for utilizing carbon substrates clustered by infection phase (MRPP, *p*=0.002), but not by mouse or time point (MRPP, *p*>0.05) (**Fig. 4A**). Examining the differential expression of these carbon utilization genes across infection phases revealed distinct metabolic patterns. Early samples expressed a broader range of substrates, marked by differential expression of D-xylose isomerase (*xylA*). Supporting this, our metabolite data denoted consumption of xylose in the infected HFD samples **(Fig. S3)**.

**Figure 4:**
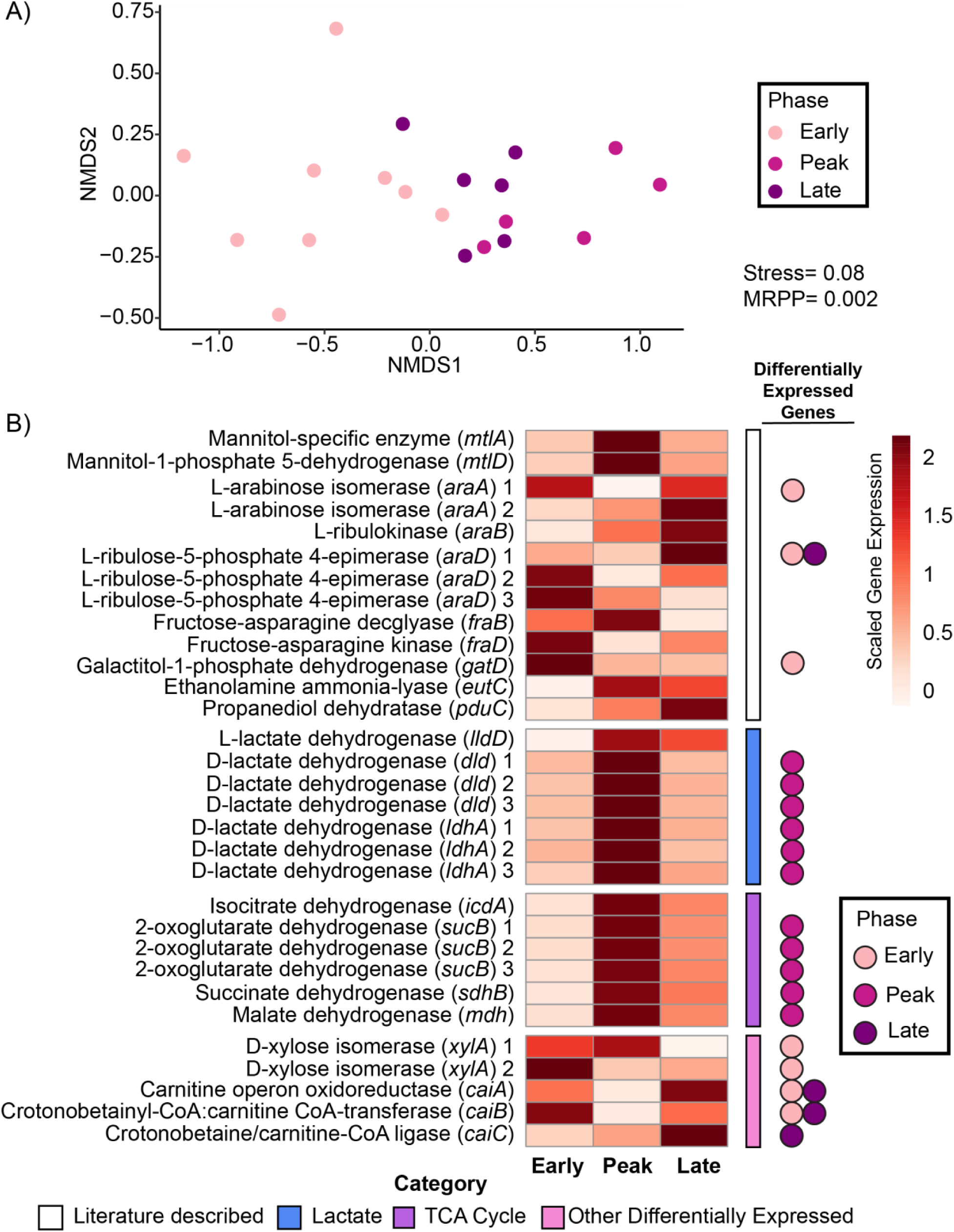
Differential expression of *Salmonella* carbon utilization genes by infection phase. A) Non-parametric multidimensional scaling (NMDS) depicts annotated *Salmonella* carbon utilization gene expression of samples colored by infection phase (early: light pink, peak: dark pink, and late: purple). Bray-Curtis dissimilarity matrix from early, peak and late samples (stress=0.08) show significant grouping of *Salmonella* carbon utilization genes by infection phase (MRPP, *p*=0.002) as found in **Data set S3**. B) Heatmap of the mean, normalized carbon utilization gene expression from our *Salmonella* pangenome shows patterns of carbon utilization by infection phase. Genes are grouped by carbon categories (Literature described carbon: white, SCFA: blue, TCA cycle: purple, Other differentially expressed carbon: bright pink). Genes that were characteristic of a specific phase (DESeq2 *padj.* >0.05) are denoted on side by colored circles (early: light pink, peak: dark pink, and late: purple) (**Data set S2**).

We also saw expression of carnitine utilization genes (*caiABCT*) (**Fig. 4B**), which were differentially expressed in late samples compared to peak samples. Carnitine has been shown to stimulate anaerobic growth of *Salmonella,* which our respiration data suggest occurs at the early and late phases (**Fig. 3**) (57, 58). Along with gene expression in late samples, we detected carnitine at day 7, the late phase of infection in HFD-infected mice **(Fig. S3)**. Further research is needed to understand the impact of these metabolisms on *Salmonella* growth and physiology, as well as the interactions with the surrounding community.

Consistent with aerobic respiration being a hallmark of peak infection, we observed simultaneous differential expression of genes for utilizing isocitrate, succinate, and malate *(icdA*, *mdh*, *sdhABCD*, *sucABCD*). It is possible that these genes were co-expressed with respiration due to roles in transforming intermediates of the tricarboxylic acid cycle, a critical component of aerobic respiration. However, it has also been shown that *Salmonella* can utilize microbially derived succinate as a substrate during aerobic respiration(21). Here the metabolite data showed less coordination with gene expression data, as succinate increased in the HFD-infected samples**(Fig. S3)**. It is possible that succinate was not used by *Salmonella* more than its microbial production, or that the metabolite was additionally host-derived, as indicated in other HFD mouse studies(59). Unraveling the complex interactions of the host-microbiome-pathogen food web are warranted for this important gut metabolite(60).

Our targeted and untargeted transcriptomic approaches revealed the significance of lactate to overall *Salmonella* energy metabolism. Of the three lactate dehydrogenase genes *Salmonella* contains (*ldhA*, *dld,* and *lldD*), the *ldhA* and *dld* genes encode an enzyme specific for the D-isomer of lactate, while the *lldD* gene encodes a protein with specificity for the L-isomer(19). It is thought that the host only produces the L-lactate isomer, while the microbial members can produce both isomers. Studies with gnotobiotic or microbiota-reduced mice have demonstrated the importance of L-lactate dehydrogenase (lldD) for *Salmonella* in utilizing host-derived lactate(19). Our targeted data show that the *lldD* gene was a core member of the transcriptome, detected across all time points but not distinguished by infection phase (**Fig. S4A**). However, our untargeted approach revealed that genes for utilizing D-lactate (*ldhA* and *dld*), likely derived from microbial production, were differentially expressed during peak infection when *Salmonella* was likely most rapidly growing based on respiration genes and ribosomal protein expression (**Fig. 4B**). Additionally, our metabolite data confirmed elevated levels of this compound at day 7 of infection relative to non-*Salmonella* inoculated mice on either diet **(Fig. S3)**, indicating production exceeding consumption during the late phase of infection. This finding suggests new cross-feeding between the microbiome and *Salmonella*.

### Non-nutritional genes expression patterns have implications on pathogenesis and horizontal gene transfer

Beyond nutritional requirements, we mined our data for other genes that were differentially expressed between phases and found categories of *Salmonella* pathogenesis genes which could be potential targets for therapeutic interventions. Volcano plots revealed gene expression patterns associated differentially expressed genes between phases (early to peak/peak to late) with the following categories: (i) not significant (5549/6473), (ii) conjugation genes (22/NA), (iii) motility genes (9/28), (iv) outer membrane genes (33/13), (v) phage-like genes (22/4), (vi) other significant genes (940/141), and (vii) hypothetical genes (127/17) (**Fig. 5**, **Data set S2**).

**Figure 5:**
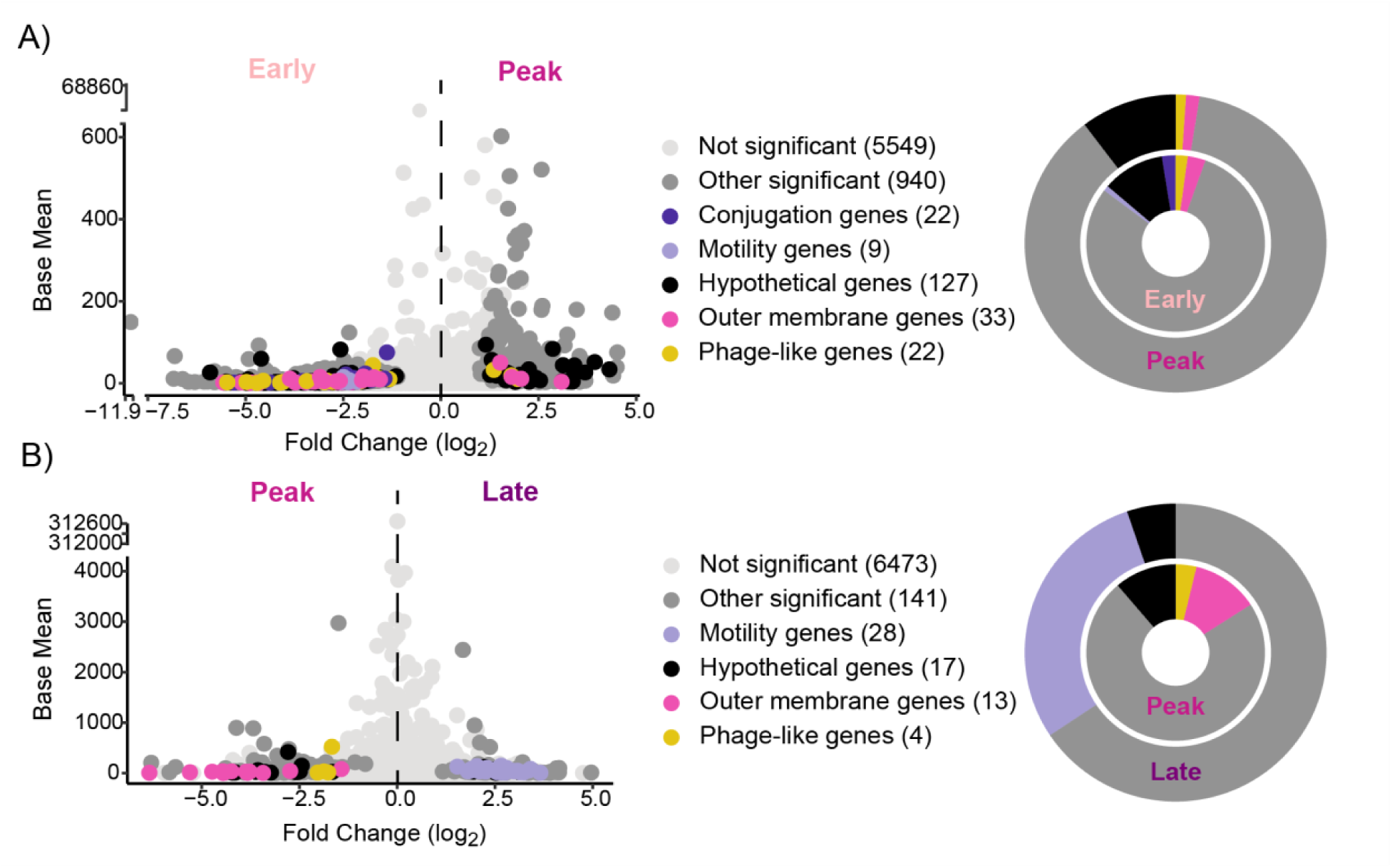
*Salmonella* pathogenesis gene expression between infection phases. Volcano plots (left) display differentially expressed genes between (A) early and peak or B) peak and late samples. Each point represents a single gene, with point color indicating significance (DESeq2, *padj.* >0.05) and annotation category. Numbers in parentheses next to each category indicate the number of genes in the *Salmonella* pangenome represented. Donut plots (right) highlight the proportion of significant genes within each category.

In early samples, we observed an upregulation of conjugation, motility, and fimbriae related genes (**Fig. 5A and Fig. S5**). Conjugation facilitates the spread of virulence genes within *Salmonella* populations, influencing pathogen evolution (61). Additionally, motility and fimbriae genes support *Salmonella* movement and adhesion, which assist interactions with colonocytes and trigger the immune response(62–65). Noteably, motility genes were also differentially expressed in the late phase compared to the peak phase. We consider it possible that *Salmonella* has enhanced chemotaxis to find nutrient sources during the late phase or for environmental entry(66, 67).

The expression of many pathogenesis genes could not be discriminated by infection phase. For instance, type III secretion protein genes (*invAG*, *sptP*, *sspH12*, *srfJ*) **(Fig. S4**, **Data set S2)** were detected and highly expressed in all infection phases. Given their presumed role in initiation of infection and the inflammatory response, it was somewhat surprising that these genes were not discriminant of early expression but instead, seemed active over the course of infection. We consider it possible that sub-populations of *Salmonella* may be infecting host cells continually or inflammation-induced envelope stress might alters the expression of these genes, explaining their chronic expression(48, 68–70).

Comparatively, peak samples differentially expressed various outer membrane associated genes (**Fig. 5B, Fig. S6**). Many of these genes were responsible for colanic acid synthesis (*wcaACDEFIJLM*). These genes protect bacteria from osmotic stress and are linked to biofilm formation(71–73). Additionally, cellulose synthase (*bcsA*), a sigma factor regulating genes controlling biofilm formation (*rpoS*) and a biofilm-dependent modulation protein were differentially expressed in the peak phase. Other biofilm-related genes (*adrA*, *csgACEFG*, *bcsBCE*, *mrlA*, *ompR*) are highly expressed in the peak and late phases. This aligns with previous findings identifying *Salmonella* luminal biofilms, where nitrate mediates two *Salmonella* populations resulting in virulent, planktonic cells and survival-adapted biofilm cells(74). Moreover, we detected differential expression of phage-like genes in both early and late infection. These prophage regions of the *Salmonella* genome carry virulence factors and are important for infection(47, 75, 76). Inflammation has been shown to boost prophage transfer between *Salmonella* species(77), but the role of phage in controlling *Salmonella* pathogenesis requires further investigation.

### Conceptual model of *Salmonella* metabolism

In conclusion, this research contributes to the development of a conceptual model illustrating how the high-fat diet background impacts *Salmonella* gene expression **(Fig. 6, Data set S2).** We show that *Salmonella* responds to a highly inflamed gut environment and tactically uses respiratory electron acceptors and carbon sources over time. For example, our findings indicate differential isomer utilization of lactate, a critical gut SCFA(19). These findings highlight the potential microbial cross-feeding as well as affirm lactate consumption across infection phases, supporting the importance of this metabolite to *Salmonella*. When possible, we supported the gene expression data with metabolite data to provide additional insights into gut metabolite transformations (**Fig. 6**).

**Figure 6:**
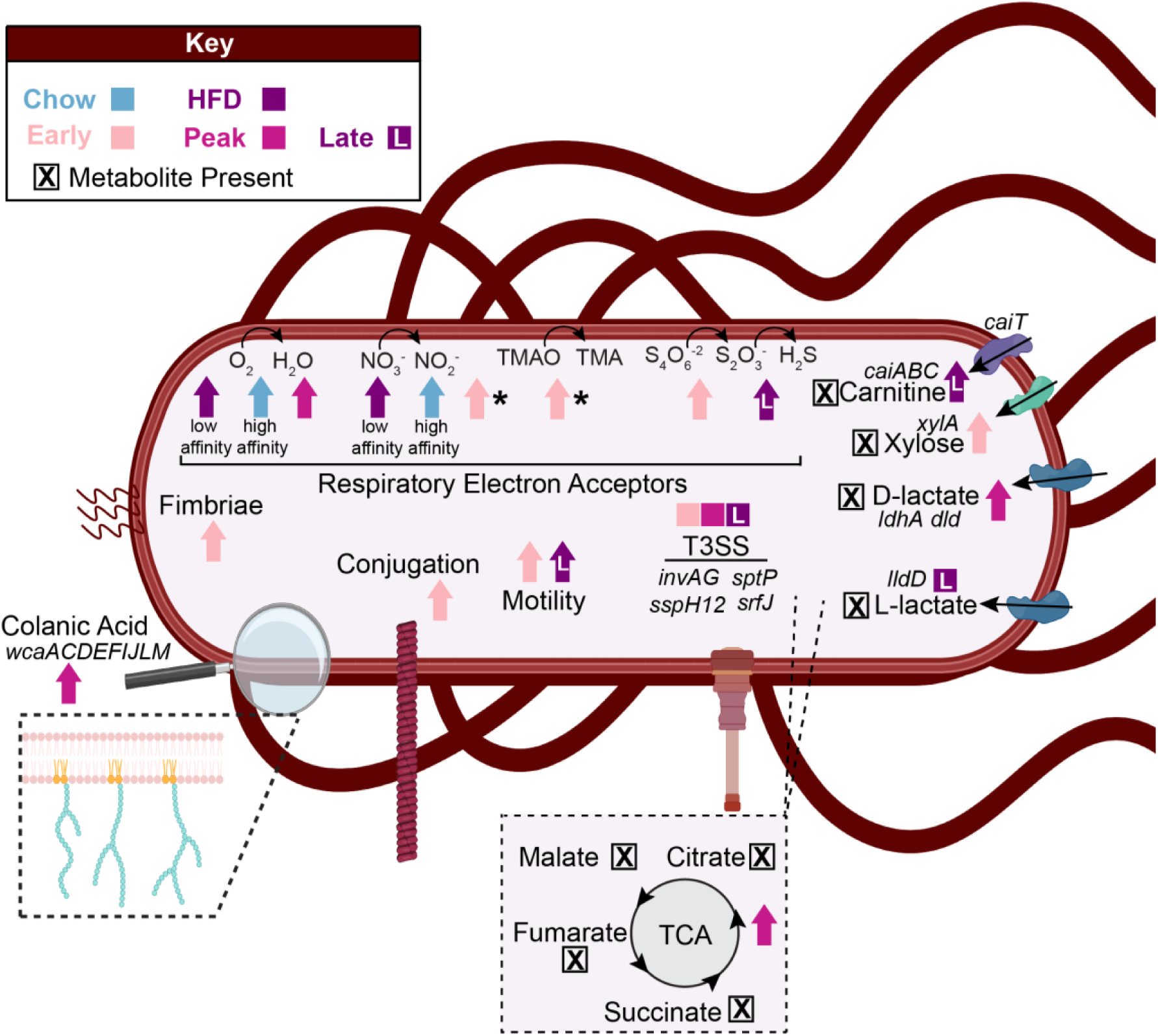
Conceptual model of findings across diet treatments and infection phases. Conceptual model summarizes our findings with regards to respiration, carbon utilization, other pathogenesis pathways, and non-differentially expressed active genes. Arrows indicate the treatment (chow: blue, high-fat diet: purple) or infection phase (early: light pink, peak: dark pink, late: purple with the letter “L”) with differential expression. Asterisk (*) next to an arrow indicates differences between DESeq2 and GeTMM, where DESeq2 was prioritized for figure creation. Boxes with an “X” show metabolite presence (**Data set S2, Data set S4**).

We also provide evidence for expression of genes related to pathogenesis, motility, and biofilms over the course of infection (**Fig. 6**). Surprisingly, virulence factor genes were not confined to the early stages when they are thought to function, but expressed across infection, indicating a heterogeneity in infection processes that were active even in a well-controlled, clonal experimental mouse model. Our discoveries benefitted from a well-established intellectual framework from years of detailed, curated pathogen physiological inquiry(11, 13, 78). This infrastructure provided a solid foundation that we could both validate and build upon. Simultaneously, our work opens new avenues for research, offering fresh perspectives and opportunities for further exploration. Moreover, this study offers a distinctive outlook on the early phases of *Salmonella* infection, providing gene expression data on days 1, 3, and 5 post-inoculation. These insights may enable research into curbing *Salmonella* proliferation during this critical period.

## Conclusion

Despite being one of the most studied microbes, knowledge of *Salmonella* metabolism and pathogenesis in relevant diet contexts and across infection phases remains limited. Here, we addressed this knowledge gap using a multi-omics approach, allowing us to examine existing theories (targeted approach) and develop new potential hypotheses (untargeted approach). In the targeted approach, we applied existing scientific knowledge to investigate specific genes previously implicated in *Salmonella* metabolism(4, 12, 13, 19–22), while the untargeted approach mines the data generated here for newly expressed functionalities discriminant of different infection phases. This work demonstrates the importance of time-dependent analysis in comprehending the finely-tuned gene expression of *Salmonella* in response to the dynamic pathobiome environment.

This research lays the foundation for understanding how *Salmonella* pathogenesis and metabolism change under realistic dietary conditions within a dynamic gut ecosystem. Unraveling the intricacies of *Salmonella* metabolism can reveal key interaction junctures with the host and surrounding microbiota. The practical implications of this study may extend to the development of targeted therapeutics designed to disrupt specific pathogenic pathways or molecules, with the aim of minimizing adverse effects on host and microbiome functionalities.

## Materials and Methods

### Strains and media

*Salmonella enterica* serovar Typhimurium strain 14028 was cultured at 37 °C in Luria-Bertani (LB) broth overnight. This culture was washed and resuspended in water for inoculation.

### Mouse experimentation and sample collection

Female CBA/J mice from The Jackson Laboratory (Bar Harbor, ME) were housed by treatment, with five mice per cage. Mice were fed either a fibrous chow diet (with 5.8% fat and 18.3% fiber, formula 7012, Teklad Diets) or a high-fat, no-fiber diet (with 36% fat and 0% fiber, formula F3282, Bio Serv) for six days before infection. Five HFD mice and 12 Chow mice were not inoculated with *Salmonella*, while the remaining mice (HFD=5, Chow=43) were inoculated with 10^9^ CFUs of *S. enterica* Typhimurium strain 14028 via oral gavage on day 0 without treatment throughout the course of infection. Mice for multi-omics analysis were selected based on fecal sample availability. High responders among chow-fed, infected mice were chosen based on *Salmonella* reaching ≥25% relative abundance at any timepoint. Noteably, unlike HFD mice, most Chow mice (n=30) are not high responders. Animal experimentation was approved by the Ohio State University Institutional Animal Care and Use Committee (IACUC; OSU 2009A0035). HFD mice were sacrificed on day 8 due to severe disease, following IACUC protocols, while Chow mice were sacrificed on day 16. Fecal samples were collected daily starting at diet transition until sacrifice (except on day −5 and −4) on autoclaved aluminum foil, transferred into labeled microcentrifuge tubes, and flash-frozen in liquid nitrogen. Samples were stored at −80°C until further processing.

### DNA and RNA extraction and sequencing

Total nucleic acid was extracted using the Quick-DNA Fecal/Soil Microbe Miniprep Kit (Zymo Research) and stored at −20 °C until amplicon sequencing could be performed. Amplicon sequencing was submitted to Argonne National Lab at the Next Generation sequencing facility, using the Nextera XT DNA Library Preparation kit (Illumina) (**Data set S1)** and the Illumina MiSeq with 2 x 251 bp paired-end reads following HMP protocols(81). PCR amplification (30 cycles) of the V4 hypervariable region of the 16S rRNA gene was conducted with universal primers 515F and 806R, with the 515F primer containing a unique barcode.

RNA was extracted using the ZymoBIOMICS DNA/RNA Miniprep Kit (Zymo Research) and stored at −80 °C until metatranscriptomic sequencing could be performed. RNA clean-up and library prep were performed using either the Ribo-Zero(TM) rRNA Removal Kit (Epicentre) with the Illumina Truseq Stranded RNA LT kit (Illumina) or Zymo-Seq RiboFree Total RNA Library Kit (Zymo Research)(**Data set S1)**. Chow samples were sequenced on the Illumina HiSeq2500 platform using 151bp paired-end reads at the Genomics Shared Resource facility at the Ohio State University. High-fat diet samples were sequenced on the NovaSEQ6000 platform on a S4 flow cell using 151bp paired-end reads at the University of Colorado-Anschutz Medical Campus at the Genomics Shared Resource Center.

### 16S rRNA Amplicon Sequencing Analysis

Data was processed using Qiime2 2019.10(82) with specific steps described here. In short, raw data fastq files were demultiplexed in Qiime 2. Then DADA2 was used for quality filtering and amplicon sequence variant (ASV) assignment. Taxonomy was determined via SILVA release 132 SSU Ref NR 99(83). Counts were filtered to ASVs with at least 10 reads in at least 5 samples. ASV feature table and taxonomic assignment are included (**Data set S1**).

### Long read sequencing and *Salmonella* pangenome generation

Genomic DNA for long read sequencing was extracted from our *Salmonella enterica* serovar Typhimurium strain 14028 isolate using the Quick-DNA Fecal/Soil Microbe Miniprep Kit (Zymo Research). Library preparation was performed using the Genomic DNA by Ligation (SQK-LSK 109) kit by Oxford Nanopore following the manufacturer’s instructions, and sequenced on the Flongle Flow Cell (R9.4.1) (Oxford Nanopore Technologies, Oxford, UK). Bases were called using Guppy (v 5.0.11), assembled using Flye (v2.8.3), and polished with long reads(84, 85). Our pangenome was created by concatenating called genes [DRAM (v1.4.0)(86)] from our highest quality *Salmonella* short read metagenome assembled genome(33) and our best long read *Salmonella* assembled genome. After filtering duplicate genes at 99% minimum sequence identity using Mmseqs2 (Release 7-4e23d)(87), our *Salmonella* pangenome is 99.99% identical to the Joint Genome Institute *Salmonella* isolate. Genes were annotated using DRAM (v1.4.0)(86).

### Metranscriptomics Data Analysis

Reads were quality trimmed and had adapters removed using bbduk.sh (v38.89)(88) and mapped to our *Salmonella* pangenome using bowtie2 (v2.4.5)(89) using flags -D 10 -R 2 -N 0 -L 22 -i S,0,2.50. Mapping files were filtered for high sequence identity (≥97%) using reformat.sh(88) and sorted by sequencing name using Samtools (v.1.9)(90). Counts were generated using htseq using flags -a 0 -t CDS -i ID --stranded=reverse (v21.0.1)(91) and normalized using DESeq2(92) or GeTMM(93) in R. For DESeq2 normalization, groups were based on infection phases (early, peak, and late). Infection phases were determined by GeTMM normalized *Salmonella* S3 ribosomal protein (*rpsC*) expression. Samples were grouped into infection phases (early, peak, and late) based on the greatest increase in *Salmonella* S3 ribosomal protein per mouse, which is the peak sample for that mouse. Any samples before the peak S3 ribosomal protein expression were the early phase, whereas any samples after were the late phase. The five samples prior to *Salmonella* infection on day −1 were used as a control for nonspecific mapping.

### Lipocalin-2 ELISA Assay

Fecal samples were homogenized in PBS containing 0.1% Tween 20 (100 mg/ml) for 20 minutes then the resulting suspension was centrifuged at 12,000 rpm for 10 minutes at 4 °C. The inflammation marker, Lipocalin-2, was measured from the resulting supernatant using the Duoset murine Lcn-2 ELISA kit (R&D Systems, Minneapolis, MN). Measuring lipocalin-2 (Lcn-2) is a tractable, sensitive marker of host inflammation(35).

### Metabolomics Sequencing and Analysis

For untargeted metabolomics, we used a 1mL solution of three solvents (water/methanol/dichloromethane,1/2/3, v/v/v) to extract metabolites from fecal samples, disrupted with a sonicator(Bioruptor®, Diagenode, Belgium). The resulting aqueous layer suspension was analyzed using Ultimiate 300 liquid chromatography coupled to Thermo Q-Exactive plus mass spectrometer (Thermo Fisher Scientific, CA, USA) coupled to a mass spectrometer with two different separation columns (reverse phase liquid chromatography and hydrophilic interaction liquid chromatography (HILIC)) for metabolome analysis. For reverse phase separation, water with 0.1% (v/v) formic acid and acetonitrile with 0.1% (v/v) formic acid were used as mobile phases. The flow rate was set at 0.3 mL/min with the gradient as follows: 2% B for 0-2 min; 2%-30% B for 4 min; 30%-50% B for 8 min, 98% B for 1.5 min, and held at 98% B for 1min, then returning into initial gradient for equilibrium for 1.5 min. For HILIC separation, ACQUITY UPLC® BEH HILIC 1.7 µm (2.1 × 150 mm) was used. Water/acetonitrile with 0.1% formic acid and 10 mM ammonium formate were prepared as solvent A (95/5, v,v) and solvent B (5/95, v,v). For gradient elution, 99% B was held for 2 min, gradually reduced to 75% B for 7 min and reduced again to 45% B for 5 min. The gradient was held at 45% B for 2 min, returned to the initial gradient and re-equilibrated for 5min. The flow rate was set at 0.3 mL/min. The quality control (QC) sample was prepared for each sample and analyzed after every 6 samples. For data processing, peak-picking and metabolome annotation were processed with MS-Dial (v.4.90)(94).

For targeted metabolomics, the short chain fatty acids were extracted as described above, and prepared using a previously published method(95). Briefly, 200 mM 3-NPH (3-nitrophenylhydrazine), 200 mM EDC (*N*-(3-dimethylaminopropyl)-*N*′-ethylcarbodiimide) and pyridine were added to extracted fecal SCFAs. Isotope-labeled SCFAs (^13^C_2_-acetic acid, ^13^C_3_-propionic acid, and ^13^C_4_ – butyric acid) were added as an internal standard before derivatization. LC-MS/MS analysis of SCFAs was conducted using the ultimate 300 liquid chromatography and Thermo Quantiva Triple Quadrupole mass spectrometer (Thermo Fisher Scientific, CA, USA). Total run time for LC was 10 min with water with 0.1% of formic acid as mobile phase A and acetonitrile with 0.1% of formic acid as mobile phase B. The gradient started with 2% B, held for 0.5 min, linearly increased up to 98% B for 8 min, and re-equilibrated in 2% B for 1.5 min. Multiple concentration of standard SCFAs (acetic acid, propionic acid, and butyric acid) were prepared alongside fecal SCFAs for quantitative analysis. The collected MS data were analyzed with Skyline(96).

### Statistical analysis

Alpha diversity metrics, richness and Shannon’s diversity, and significance values were calculated in R using the vegan package (v2.5-7)(97). To compare carbon utilization expression patterns among samples, Bray-Curtis dissimilarity was calculated using 406 *Salmonella* carbon utilization genes annotated by DRAM (**Dataset S3**). Annotation calls for CAZymes, central carbon, hydrocarbon, and pyruvate metabolism were selected as well as carbon associated genes from our *Salmonella* DRAM module. Non-parametric multidimensional scaling (NMDS) plots were created using R (ggplot2 package v3.3.5, and the vegan package (v2.5-7) for visualization and non-parametric-fit quality was determined by stress value(97–99). Significance of infection phase carbon utilization expression differences was determined by analysis of similarity (ANOSIM) and multiple response permutation procedure (MRPP)(97). Heatmaps were generated with GeTMM normalize expression scaled by gene using the R package pheatmap (v1.0.12)(100).

## Supporting information

Description of Supplemental files

Data set S1

Data set S2

Data set S3

Data set S4

## Data Availability

All data files and R scripts to generate figures are available in Github at https://github.com/Kokkinias/HFDtimeseries. All *Salmonella* MAGs and raw data is deposited at the National Center for Biotechnology Information (NCBI) under accession number PRJNA348350. The gene delineated *Salmonella* pangenome is available in Zenodo at DOI: <u>10.5281/zenodo.10479610</u>.

## Acknowledgements

This work was supported by NIH NIAID R01AI143288 awarded to VHW, BMMA, and KCW. Additional support came from NIH Predoctoral Training grants T32AI162691 (KK) and T32GM132057 (IL). The funders had no role in the study design, data collection, interpretation, or the decision to submit for publication. The microbial annotation tool, DRAM (86), received support from the DOE Office of Science, Office of Biological and Environmental Research (BER), grant no. DE-SC0021350.

We acknowledge Erin Boulanger and Andrew Schwieters for their assistance with mouse experiments. Special thanks to the integral work of Tyson Claffey and Richard Wolfe from the Colorado State University server management group, Sandy Shew for managing computing resources from The Ohio State University Unity cluster, and the Campus Chemical Instrument Center – Mass spectrometry & Proteomics at Ohio State University Comprehensive Cancer Center (OSU CCC) under the guidance of Dr. Pearlly Yan for supporting our use of metabolomics instruments.

ASD, IL, and MS conducted mouse experiments and collected fecal and cecal samples. RK and RAD performed DNA and RNA extractions, ensuring quality control. KK, YK, IL, and MS analyzed and interpreted sequencing and metabolomic data. ASD quantified lipocalin-2 using ELISA. YK and VHW managed LC-MS/MS sample preparation and data collection for metabolomics. ASD and BMMA provided expertise on *Salmonella* physiology and experimentation. MAB and KCW provided guidance on microbial metabolism and sequencing best practices. KK and KCW led the manuscript writing, with all co-authors contributing edits. The authors declare no competing interests.

---

**Figure S1:**
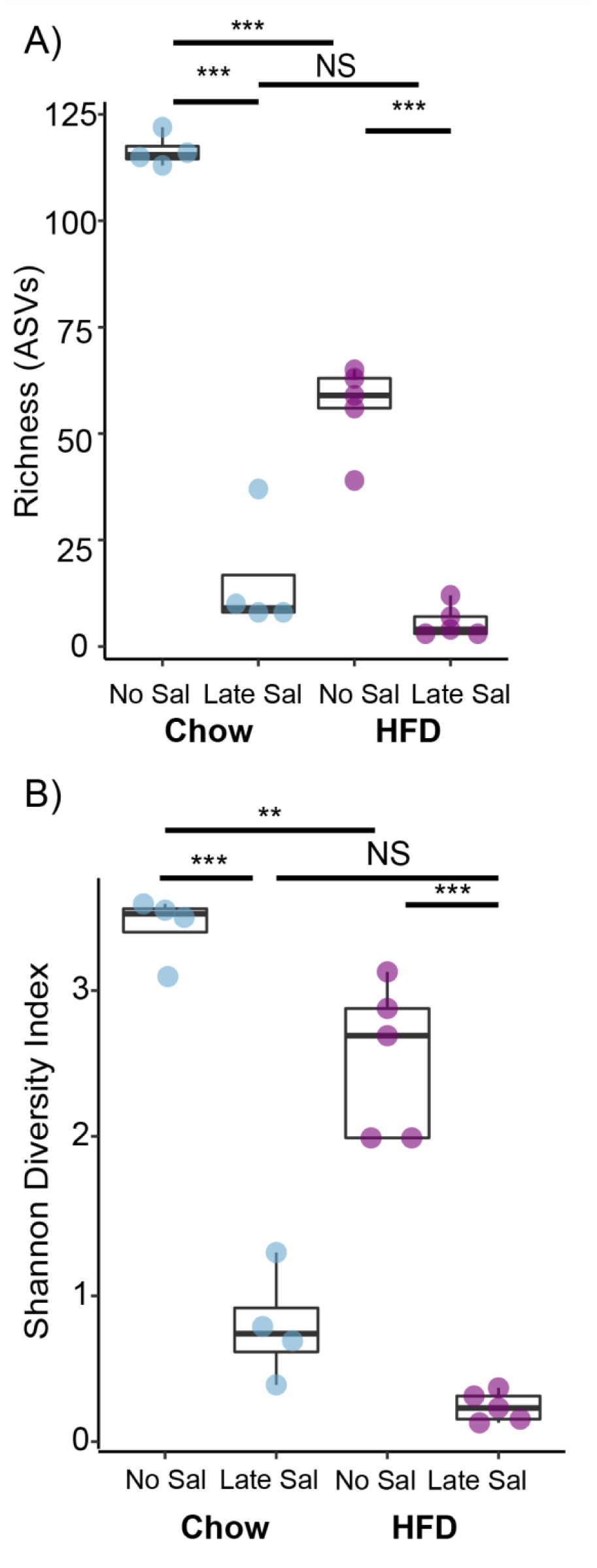
Microbial Diversity between Chow and HFD mice. A) Richness and B) Shannon’s diversity metrics between chow (Chow) and high-fat diet (HFD) mice fecal samples prior to infection (day −1) and during late infection (day 11 or day 8, respectively). Asterisks indicate statistical significance where ** is a *p* value of <0.01 and *** is a *p* value of <0.001. NS indicates that there is no statistically significant difference.

**Figure S2:**
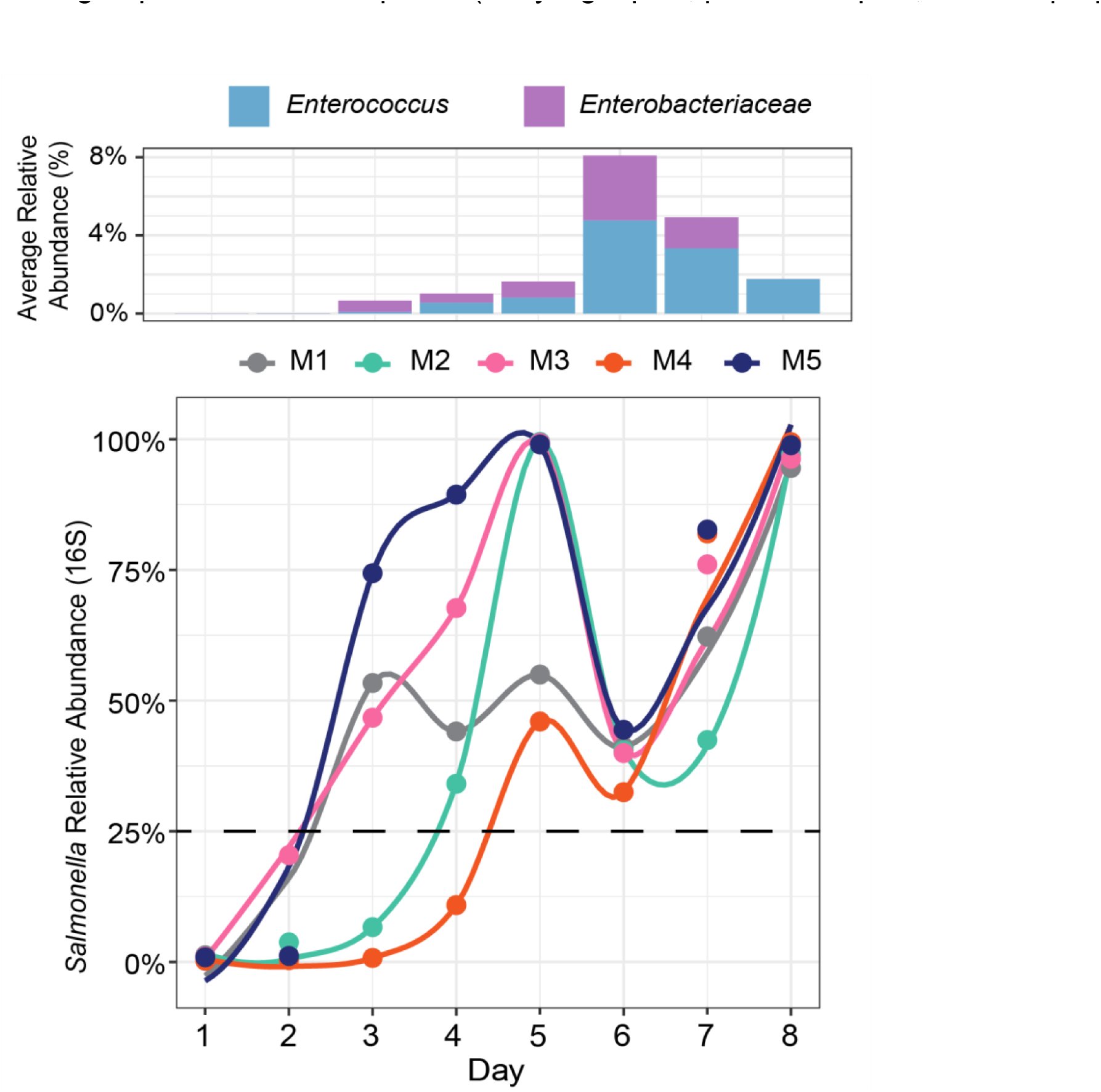
16S *Salmonella* relative abundance per mouse over time. The line plot (bottom) shows 16S rRNA gene relative abundance of *Salmonella* per mouse over time, and the bar chart (top) shows the average relative abundance of one *Enterococcus* amplicon sequencing variant (ASV) and one Enterobacteriaceae ASV over time.

**Figure S3:**
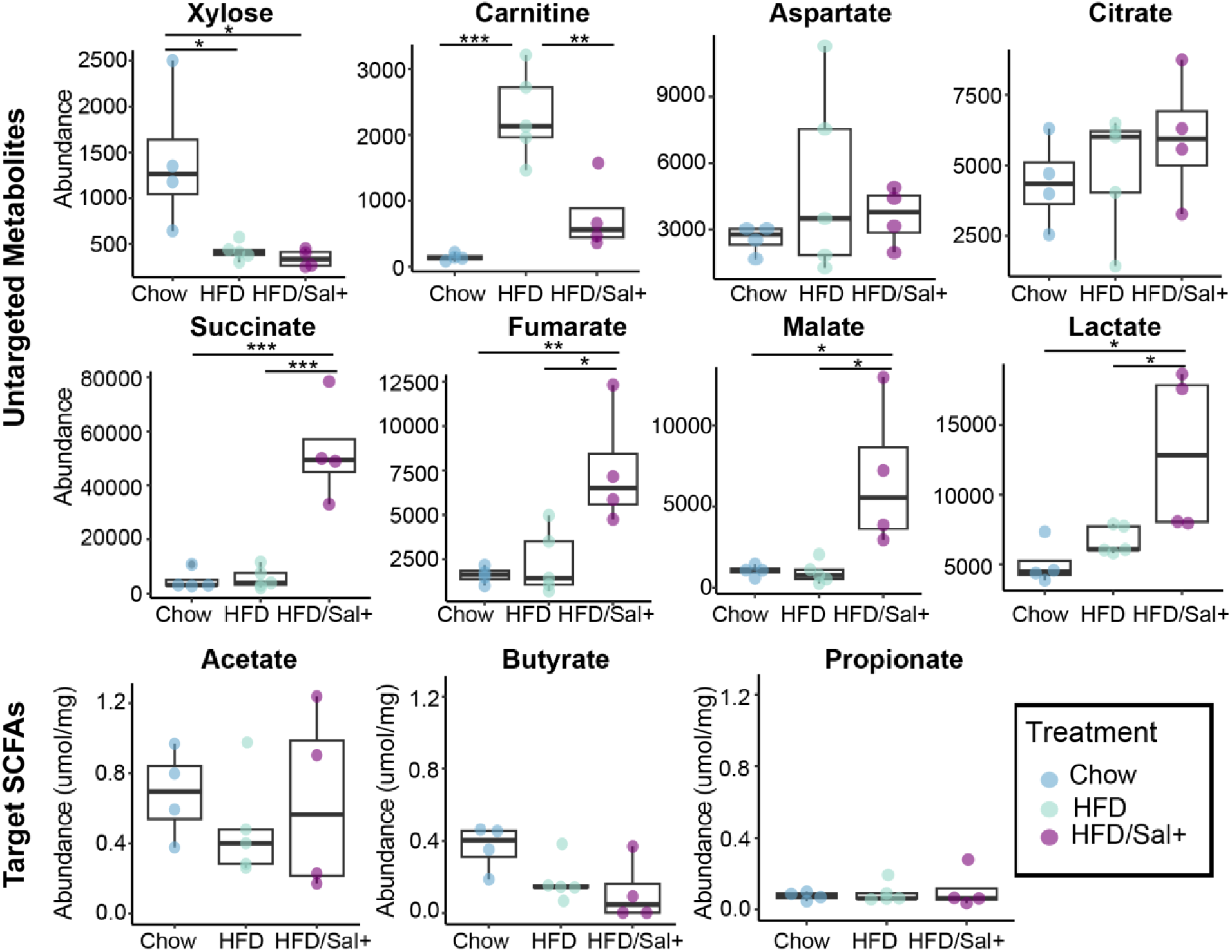
Metabolite abundance during late infection phase across treatments. Boxplots depict untargeted (xylose= C00181, carnitine= C00487, aspartate=C00049, citrate= C02226, succinate= C00042, fumarate= C00122, malate= C00497, lactate= C00256) and targeted (acetate, butyrate, propionate) metabolite abundance in fecal samples of uninfected chow (Chow, blue), uninfected high-fat diet (HFD, light green), and infected high-fat diet (HFD/Sal+, purple) mice on day 7. Asterisks indicate statistical significance where * is a *p* value of <0.05, ** is a *p* value of <0.01, and *** is a *p* value of <0.001.

**Figure S4:**
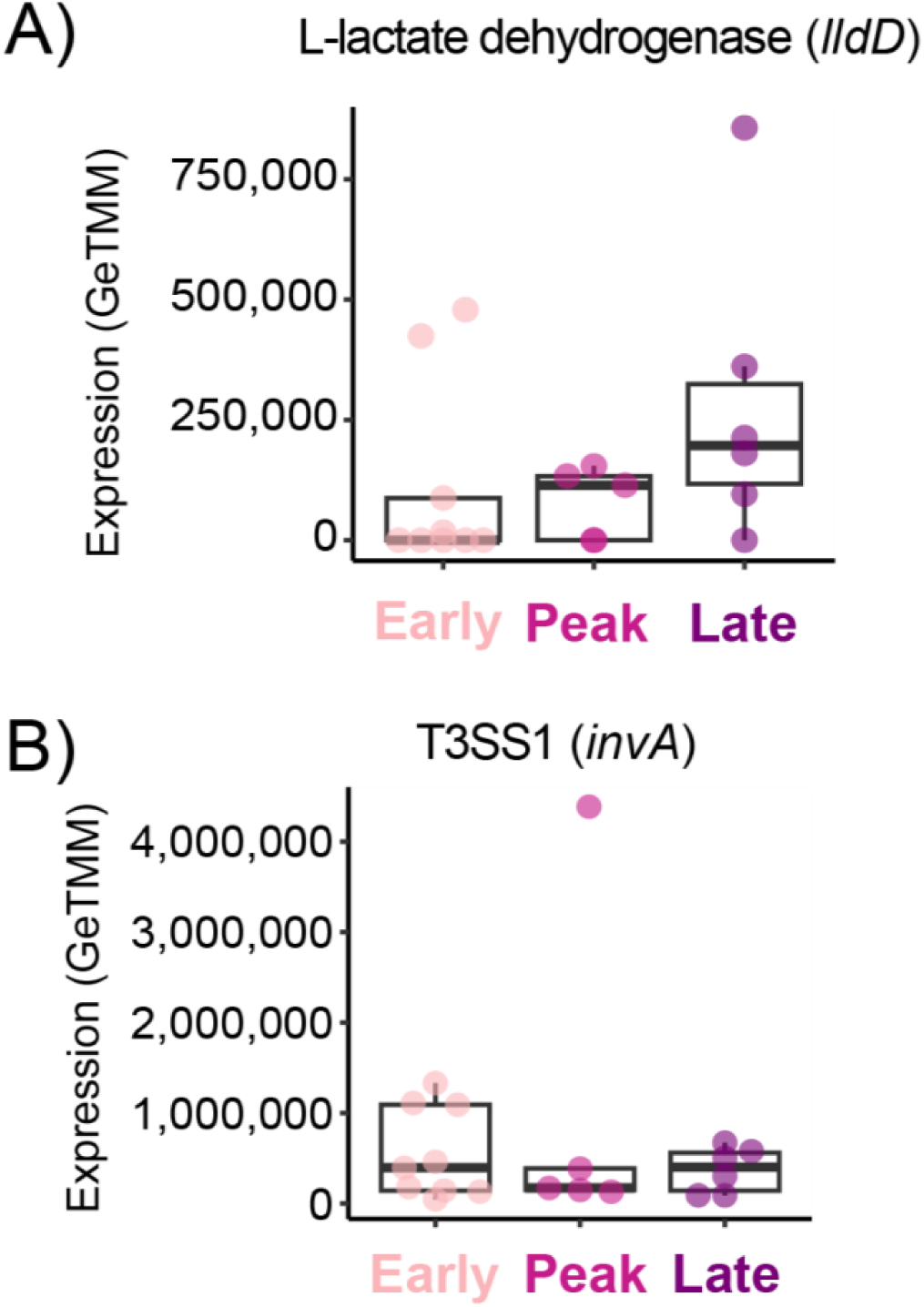
Other active, non-differentially expressed genes. Boxplots show normalized gene expression of A) L-lactate dehydrogenase (*lldD*) and B) a type III secretion protein (*invA*) across infection phases.

**Figure S5:**
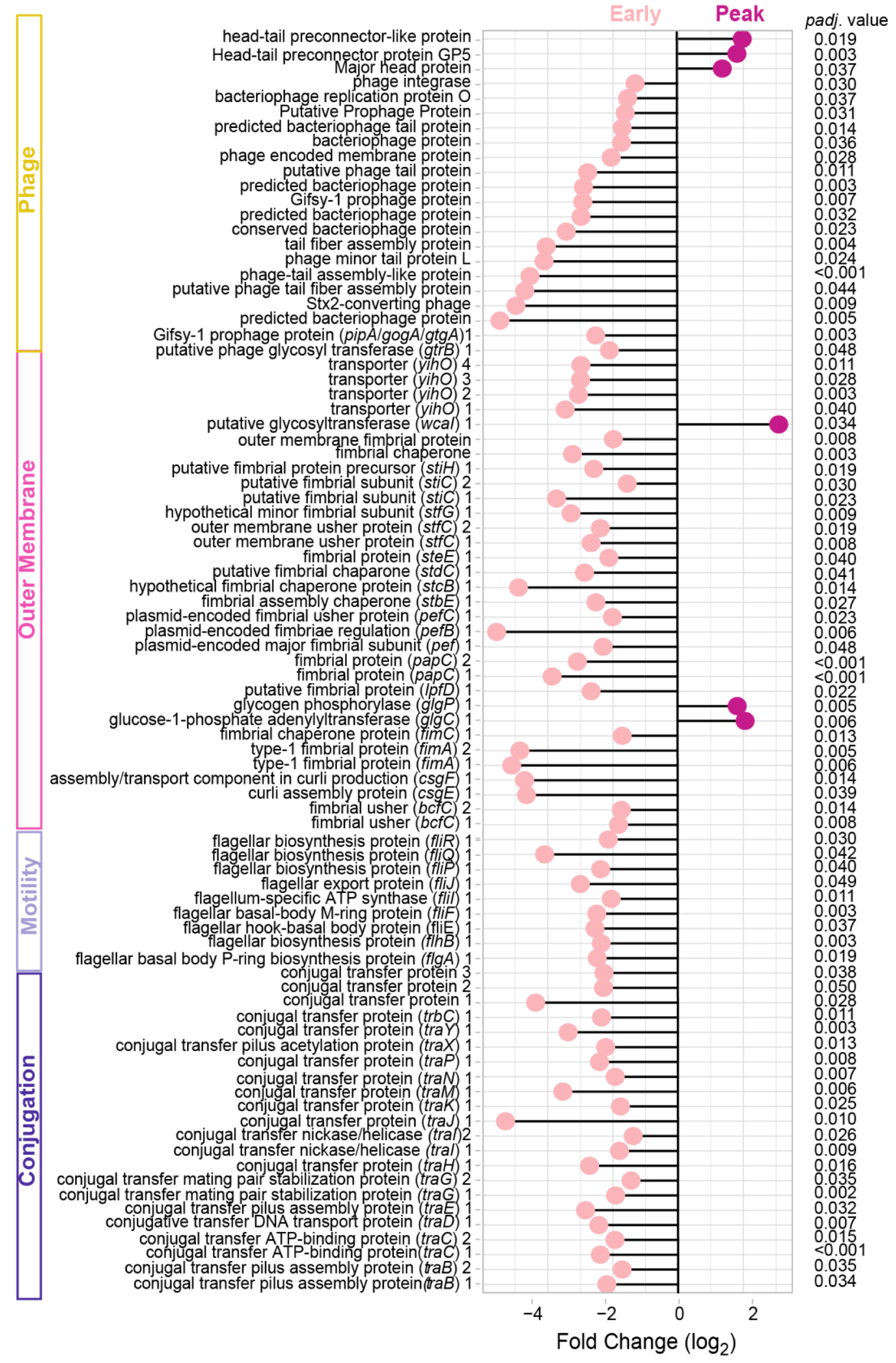
Differential expression of pathogenesis genes between early and peak phases. Lollipop plot of fold change (log_2_) of differentially expressed (DESeq2, *padj.* >0.05) pathogenesis pathway genes between early (light pink) and peak (dark pink) infection phases. Genes are ordered by gene categories: conjugation (purple), motility (light purple), outer membrane (pink), and phage-like genes (yellow). Adjusted *p* values are listed in the *padj*. value column.

**Figure S6:**
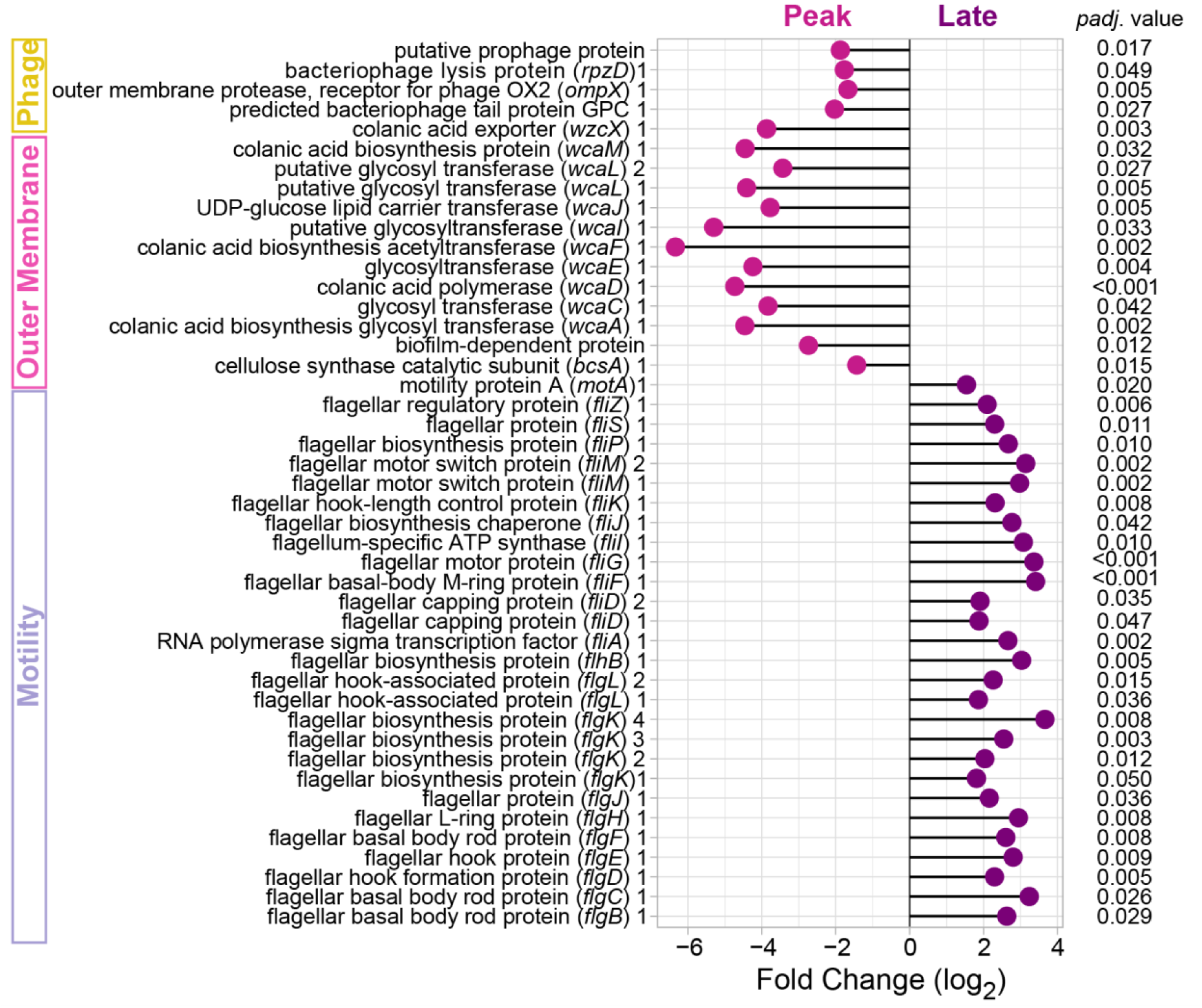
Differential expression of pathogenesis genes between peak and late phase. Lollipop plot of fold change (log_2_) of differentially expressed (DESeq2, *padj.* >0.05) pathogenesis pathway genes between peak (dark pink) and late (purple) infection phases. Genes are ordered by gene categories: motility (light purple), outer membrane (pink), and phage-like genes (yellow). Adjusted *p* values are listed in the *padj*. value column.

