## Supplementary material for "Time resolved multi-omics reveals diverse metabolic strategies of *Salmonella* during diet-induced inflammation": Description of Supplemental files

Supplemental Data File Descriptions

Data set S1 contains metadata about the multi-omics performed, ASV feature count table, SILVA taxonomy (132-99-515-806), diversity metrics, and lipocalin-2 concentrations (ng/g of feces). 

Data set S2 contains raw, unfiltered mapping to the Salmonella pangenome, gene length and strandedness information, GeTMM normalized gene counts, DESeq2 differentially expressed genes by each phase (early, peak, post), DRAM (v1.4.0) annotations, and a list of genes depicted in figure 6. 

Data set S3 contains a GeTMM normalized Salmonella carbon gene expression matrix, annotations of the Salmonella carbon gene from CAZyme and DRAM module step form, and the DRAM module step form (https://github.com/WrightonLabCSU/DRAM). 

Data set S4 contains abundance and annotations of untargeted metabolites and abundance of targeted short-chain fatty acids (acetate, propionate, and butyrate).
